# Accurate and simple FXN-GAA repeats (Friedreich ataxia loci) estimation by long read targeted sequencing

**DOI:** 10.1101/2022.02.24.481841

**Authors:** Pooja Sharma, Bharathram Uppili, Istaq Ahmad, Shweta Sahni, A Vivekananda, Mohammed Faruq

**Affiliations:** Genomics and Molecular Medicine, CSIR-Institute of Genomics and Integrative Biology (CSIR -IGIB), Mall Road, Delhi 110007, India; Academy for Scientific and Innovative Research, Ghaziabad, 201002, India; Neurology Department, Neuroscience Centre, All India Institute of Medical Sciences, New Delhi 110029, India

**Keywords:** Friedreich ataxia, Long read sequencing, Trinucleotide repeats, GAA repeats, FRDA, NGS

## Abstract

Friedreich ataxia, an autosomal recessive disorder is caused by tandem GAA nucleotide repeats expansion in intron 1 of the FXN (frataxin gene). The GAA repeats above 66 in length are considered as pathogenic and commonly occurring repeats are 600-1200. Clinically the spectrum of the features is confined mainly to the neurological tissue, however, cardiomyopathy and diabetes mellitus has been observed in 60% and 30% of the subjects. The accurate detection of GAA repeats count is of utmost importance for clinical genetic correlation as no study has attempted an approach which is of high throughput nature and defines the sequence of GAA repeats. Largely, the method for detection of GAA repeats so far is either conventional PCR based screening and southern blot which is the gold standard method. We describe for the first time a method of long read sequencing wherein we utilized approach of long range targeted amplification of FXN-GAA repeats and sequencing on oxford MinION platform. We were able to achieve the successful amplification of GAA repeats ranging from 180-1200 at 250x coverage. The total throughput achievable for 96 samples can be less than 24 hours on one flow cell as per our protocol and is scalable and deployable at clinical day to sequencing.

## INTRODUCTION

Friedreich ataxia (FRDA) is an autosomal recessive disorder that affects neurological (spinal cord, sensory nerves and cerebellum) and extra-neuronal systems (heart and pancreas). Disease develops more often at below 25 years of age. The causal mutation is a hyperexpansion of GAA repeats within intron 1 of the FXN gene on chr9q21.11. The GAA repeat length is variable varies from in healthy subjects and is observed between 5-33 in healthy subjects, while alleles greater than 66 to 1300 repeats represent full length pathogenic alleles (1,2). The commonest GAA repeat length observed in patients is ∼600-1200. The cardinal clinical symptoms and signs of FRDA include are gait ataxia, dysarthria, peripheral neuropathy, kyphoscolisosis, pes cavus, however, only ∼60% of the patients show hypertrophic cardiomyopathy and nearly one third of patients manifest diabetes mellitus. Nearly 25% of all FRDA cases do not subscribe to the essential diagnostic criterion for age of onset before 25 years as defined by Harding (3), and are referred to as the Late Onset FRDA (LOFA) (4). Atypical features in LOFA are defined by clinically milder disease phenotypes with a much slower disease progression, attributed to the size of the expanded repeat on GAA1 (5, 6).

The method of detection of FXN-GAA repeats is through conventional PCR, coupled with Triplet repeat-primed PCR which is helpful in screening; however, Southern blot is the gold standard method to detect the GAA length. The determination of precise and accurate GAA-length holds considerable great importance because the observed clinical variability in FRDA is largely unexplained at large. Some of the pertinent clinical and technical issues which warrants alternate and advanced tools to detect not only the GAA repeat length but also the repeat the interruption are, i) the determination of sequence configuration of GAA track would allow its correct interpretation if its purity of it can be ascertained vis-a-vis against the interrupted sequences, ii) the GAA sequence determination would further allow in understanding of the genotype-phenotype correlation, iii) it would allow us to decipher the relation between late onset Friedreich ataxia (LOFA) and interrupted GAA sequence, iv) southern blot is a low throughput technique for rapid and determination of GAA length. Sequence variation within the GAA repeat region has been reported to affect the phenotypic variability in FRDA, especially in cases with late onset of disease (7). The aforementioned issues underscore the need for development and deployment of a novel approach to determine GAA length via next generation sequencing.

While short read sequencing for detection of repeat mediated disorders has already progressed much, long read sequencing as offered by the Oxford Nanopore technology (ONT) offers many unprecedented advantages over the former. Besides their rapidly shrinking costs and improved read quality, long read sequencing platforms provide additional maneuvering with respect to the bioinformatics analysis pipelines. This is of special significance in diseases attributable to complex expanded alleles with multiple repeat interruptions. Long read sequencing or 3rd generation sequencing technologies are a promising toold for querying the human genome for the assessment of long tracts of tandem nucleotide repeat expansion and over the last few years, several novel genes have been identified through long read sequencing technologies, i.e SMRT and oxford nanopore based platforms. LRSeq has lead to the discovery of various new tandem nucleotide repeats (TNR) mutations which would otherwise not be detectable through short read sequencing methods. The discovery of these TNRs has opened up new avenues for solving the pathology of challenging neurological disorders, such as e.g. BAFME, NOTCH2NLC etc (8, 9).

The shortcomings of the existing approaches along with the unparalleled advantages offered by the ONT based long read sequencing makes it imperative to deploy this It is very important now to use the same technology for accurate detection of TNR sequences for known disease loci. and Efforts made in this direction in the past have been successful for FMR1-CGG loci, HD-CAG etc.

To the best of our knowledge, no study has so far reported the accurate GAA length determination using a long read sequencing platforms and the present study reports a comprehensive and simple protocol used for accurate determination of GAA repeats using nanopore technology. We were successful in estimating the length estimation of GAA repeat regions within the range of 200-1000 repeats.

## METHODOLOGY

The subjects included in the present study were recruited from our repository of DNA samples at the CSIR-Institute of Genomics and Integrative Biology. Genomic DNA was extracted using salting out method^(1)^. For the initial phase, we screened the samples for the prevalent known ataxia subtypes (SCA1, SCA2, SCA3, SCA6, SCA7, SCA12, SCA17 and FRDA) using fluorescently labeled primers.^[2]^ The PCR product size (fragment length of each of the loci) were analyzed by capillary electrophoresis on ABI 3730XL DNA Analyzer (Thermo Fishers Scientific) and size estimation through Genescan software v 4.0. For FRDA screening GAA expansion detection was performed through triplet repeat primed PCR.^[3]^ *FXN*-GAA numbers were estimated by Long Range PCR using primers (FP-^5’^GGAGGGATCCGTCTGGGCAAAGG^3’^; RP-^5’^CAATCCAGGACAGTCAGGGCTTT^3’^) with KOD plus enzyme (Kit No 201 200U, 200 Reaction TOYOBO Com. Ltd Japan).^[4]^ Long Range PCR was performed with initial denaturation at 94°C for 3 minutes. Two step PCR of 20 seconds denaturation at 94°C and 2.30 minutes annealing and extension at 68°C in addition with 15 seconds increase per cycle for total 22 cycles. The final extension was carried out at 68°C for 10 minutes. PCR amplicons were checked on 1% agarose gel with 1kb DNA ladder (Promega) and stored at 4°C. The amplified products were purified using ampure beads (beckman coulter) in 1.6x ratio by volume of the PCR product.

### Library prepration and sequencing

The purified amplicons with a length of 1.4kb to 4.5kb were sequenced by using Oxford Nanopore Sequencing platform. Around 400ng of purified long range amplicons were taken as input for sequencing. The library was prepared following Nanopore Native barcoding genomic DNA with EXP-NBD196 and SQK-LSK109 protocol according to the latest sequencing version: NBE_9121_v109_revF. The DNA amplicons were end repaired and dA-tailed using NEBNext FFPE DNA Repair Mix (New England Biolabs (NEB), Ipswich, USA and NEBNext End repair/dA-tailing (NEB). Native barcodes were ligated to the end repaired DNA with NEB Blunt/TA Ligase Master Mix (SQK-LSK109). Adaptor ligation to the pooled barcoded library was performed using NEBNext® Quick Ligation module ((NEB E6057). The final library was purified with ampure beads in 0.4x ratio of the reaction volume and Oxford Nanopore short fragment buffer (SFB-SQK-LSK109). The prepared library of around 70ng was sequenced using R9.4.1 flow cell on MinION Mk1C for around 16 hours.

### Sequence Analysis

Using Guppy basecaller (version 3.5.2) the raw fast5 files were demultiplexed into different fastq files. The alignment was performed with minimap2 3 (v 2.1) with FXN gene sequence (GRCh37 - chr9:71650479-71693993) as the reference (13, 14). The bam file is sorted and indexed using samtools 2 (v 1.9). For finding the repeat expansion inside the sample we have used STRique (14). Initially, Indexing of the fast5 was performed to tabulate all the fast5 reads from all the samples, later using the HMM model as given by the STRique tool (r9_4_450bps.model) the repeat count per each read for all the samples had been calculated. Using ggplot2, dplyr packages from R the data has been visualized.

## RESULTS and DISCUSSION

### GAA repeat amplification through long range-PCR

The long range amplification using flanking primer sequence has amplified sample specific PCR products of varying length ranging from 1,500 base pair - 4000 base pairs (Fig.1). The PCR products of length ∼1.5 kbp are expected to carry small normal GAA repeats (7-20) while fragments of 2.2 kb were expected to carry ∼200 GAA, likewise, fragments of size ∼4kb corresponds to GAA ∼1000 repeats. The long range amplification of the GAA flanking region was efficient in capturing all the expected length of abnormal GAA repeats.

**Figure 1:**
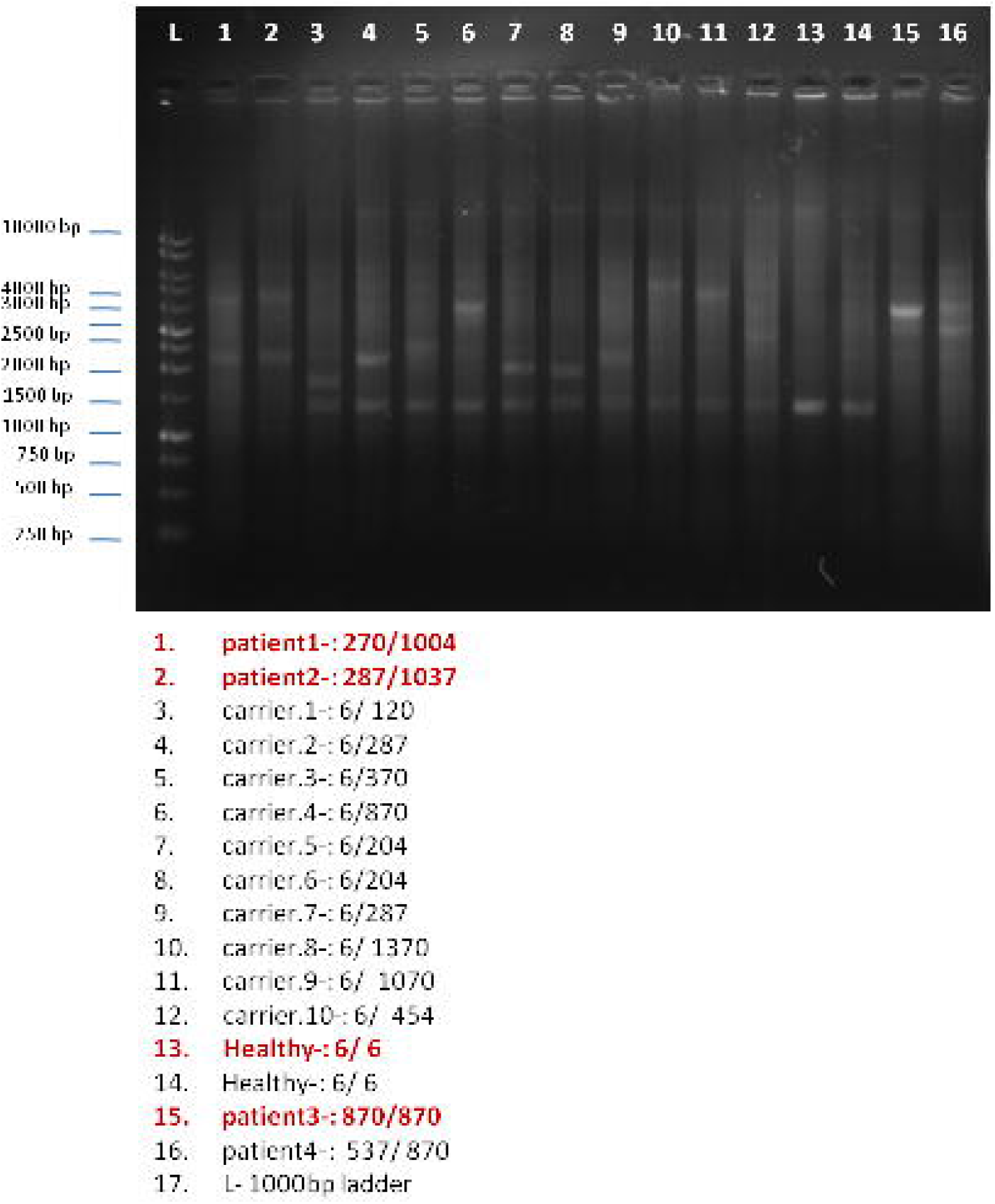
Long range-PCR base detection of GAA repeats and flanking region

### Targeted GAA flanking region sequencing on MinION flow cell

We sequenced four representative PCR products from four subjects (patient1, patient2, patient3 and a healthy subject) with expected GAA (allele1/allele2) length of 200/1000, 300/1000, 800/800 and 6/6 respectively. We obtained a total of 140k reads for four samples (average 23k reads/sample that mapped to the target region) with mean coverage ∼260x (165-350X). The data QC showed the max sequenced reads of 4kb length with some non-specificity off target longer products, the percent GC content of 42-45 and read mapping percentage ranging from 59% to 66%. (Fig.2)

**Figure 2:**
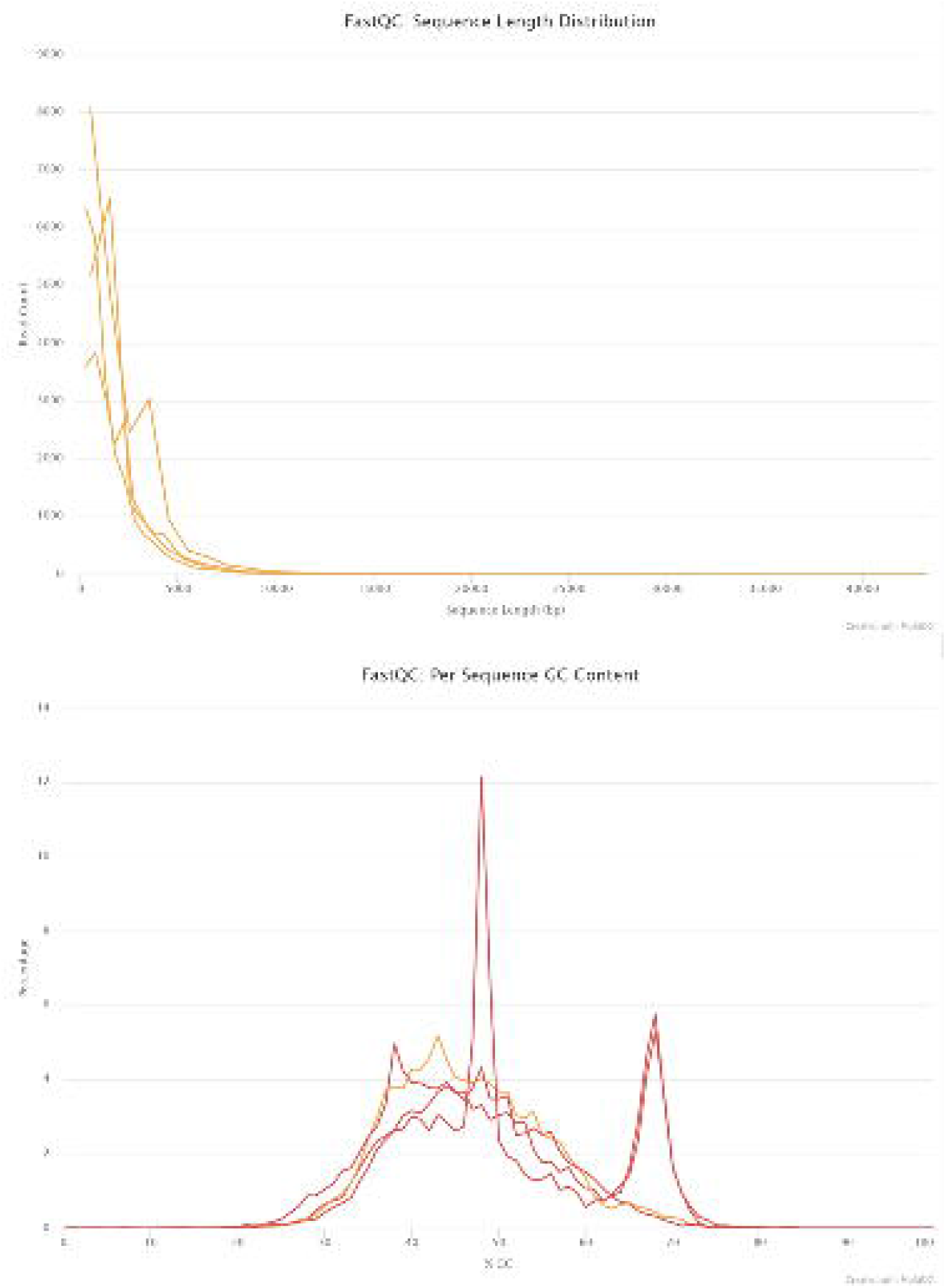
Quality parameters of the reads obtained through nanopore sequencing showing read length and % GC content.

### GAA repeat sequence determination

Using STRique, the length estimation of GAA showed for patient-1, biallelic expansions mutations with highest peak at 250 bp (allele1) and 1070 (allele2); for patient-2, allele1/allele2 (250/1100) and patient 3 (525/830). (Fig.3). the squiggle (series of current pattern in GAA repeat region was detected along with flanking non-repetitive region (fig.4).

**Figure 3:**
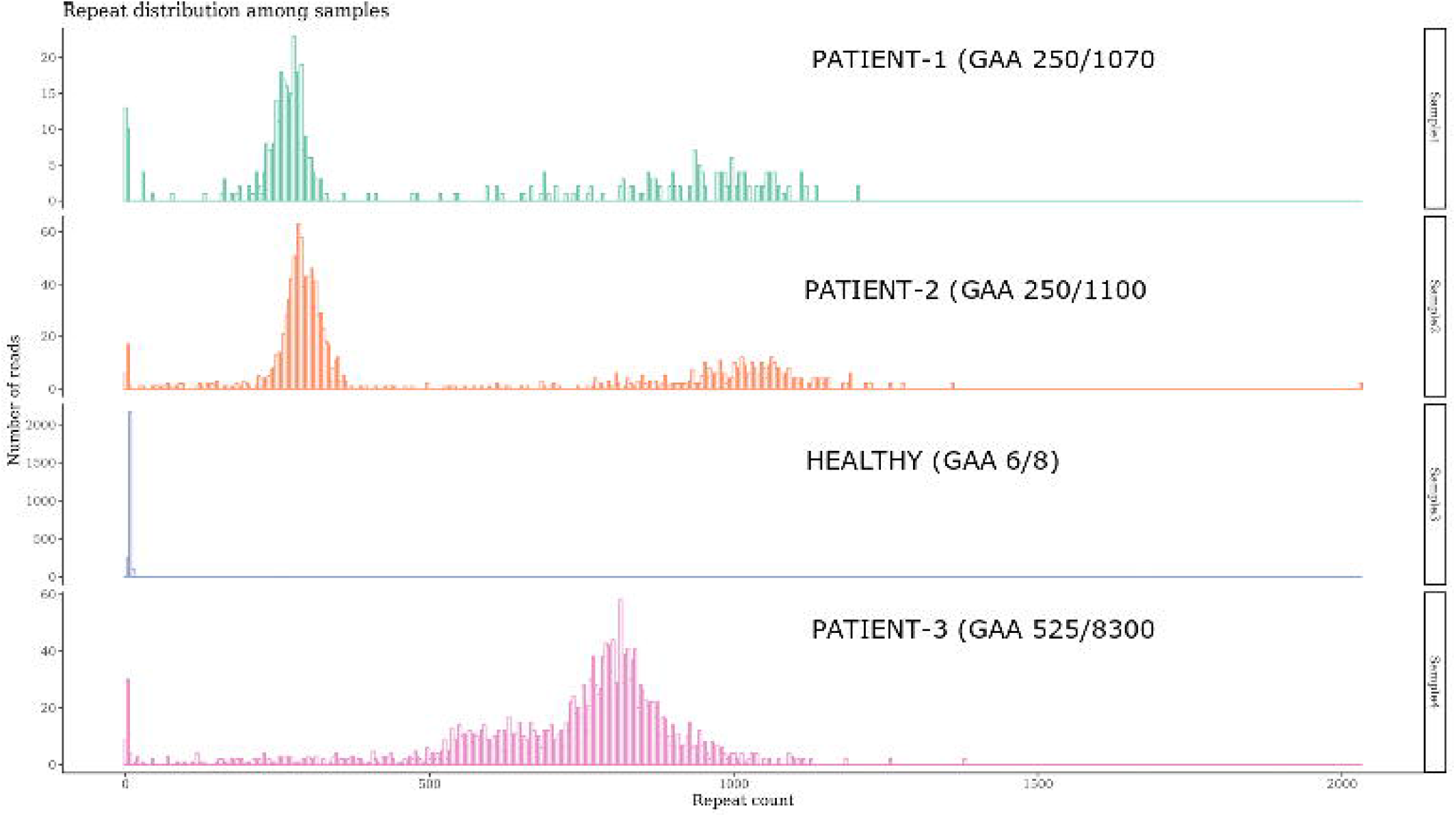
Histogram showing length wise GAA repeat distribution across four samples. The variability in length is indicative of somatic mosaicism

**Figure 4:**
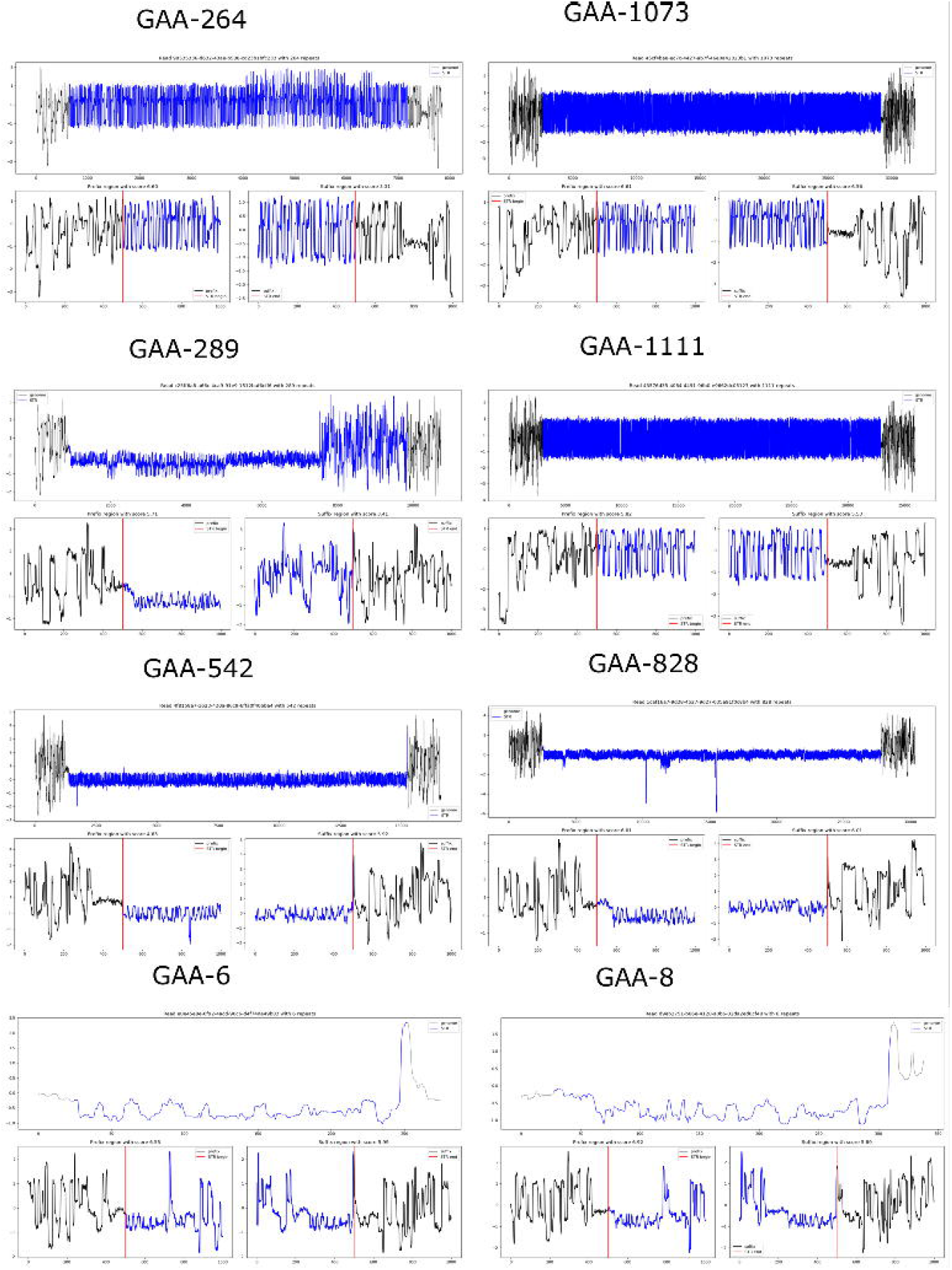
Current series pattern of representative reads in the form of squiggle for each of the samples.

### Conclusion

Despite their important role in determining the clinical variability in FRDA, GAA repeat structure investigation has so far precluded large-scale screening. Long-read sequencing with its promising ability for identification of causative repeat expansions, discernment of repeat configuration, repeat length estimation and detection of base modifications, is aptly positioned to overcome these challenges. For the first time we demonstrate the ability of long read ONT sequencing in accurately determining the size of an expanded intronic region of upto 1000 GAA repeats in FRDA patients.

## Acknowledgement

We acknowledge the funding from CSIR-IGIB, OLP1120 and ICMR funded project GAP240. We thank sincerely to the patients and the families for their participation.

## Notes

### Competing Interest Statement

The authors have declared no competing interest.

